# Mechano-Initiated PIEZO1-VEGFR2 Interaction Governs CD34^+^ Cell Differentiation and Repair in Arteriovenous Fistula

**DOI:** 10.64898/2026.06.08.731007

**Authors:** Pengwei Zhu, Yakui Wu, Linlin Lu, Tao Huang, Ruihan Chen, Yanhua Hu, Liujun Jiang, Xuliang Wang, Qingbo Xu, Jiang-Yun Luo, Xiaosheng Hu

**Author notes:** **Addresses for Correspondence:** Jiang-Yun Luo, PhD, Department of Cardiology, the First Affiliated Hospital, Zhejiang University School of Medicine, 79 Qingchun Road, Hangzhou 310003, Hangzhou, China, Or Xiaosheng Hu, MD, Department of Cardiology, the First Affiliated Hospital, Zhejiang University School of Medicine, 79 Qingchun Road, Hangzhou 310003, Hangzhou, China.

## Abstract

**Background:** Endothelial cell (EC) injury induced by disturbed flow drives neointimal hyperplasia in arteriovenous fistulas (AVFs), where CD34^+^ cell-mediated repair may be involved. PIEZO1 and VEGFR2 are important mechanosensors with critical role in maintaining endothelial function. However, whether PIEZO1 interacts with VEGFR2 during CD34^+^ cell differentiation to orchestrate the vascular repair remains unknown.

**Methods:** AVF model was established in several mouse strains. Single cell RNA sequencing was performed for human and mouse samples. *Cd34*-CreERT2; R26-tdTomato; *Piezo1*^flox/flox^ mice were used to investigate the effect of *Piezo1* deletion on endothelial repair in AVFs. CD34-high human umbilical vein ECs (CD34^high^ HUVECs) was sorted and exposed to different flow patterns to determine the role of shear stress in CD34^high^ cell differentiation. Co-immunoprecipitation, proximal ligation assay and complementary approaches were performed to delineate mechanotransduction initiated by PIEZO1-VEGFR2 interaction.

**Results:** Single cell RNA sequencing and immunostaining showed abundant CD34^high^ cells in the vessel wall of AVFs in humans and animal models. Exposure of CD34^high^ HUVECs to different flow patterns showed that laminar shear stress downregulated CD34 while upregulating VE-cadherin and claudin-5 expression. In contrast, oscillatory flow produced the opposite effects, indicating impaired endothelial maturation. PIEZO1 knockdown in CD34^high^ HUVECs attenuated shear stress-induced endothelial marker expression. In Cd34 conditional Piezo1 knockout mouse model of AVF, we observed decreased number of CD34-derived cells, more compact cellular arrangement, and attenuated neointimal hyperplasia. Mechanistically, we found PIEZO1 interacts with VEGFR2, thereby mediating the distinct effects of laminar and oscillatory shear stress on AKT-FoxO1 axis, which critically regulates endothelial marker expression. Furthermore, pharmacological activation of AKT signaling in AVF mouse model enhanced CD34^+^ cell-mediated endothelial repair and attenuated neointimal hyperplasia.

**Conclusion:** PIEZO1-VEGFR2 complex-mediated mechanotransduction plays a key role in regulating CD34^+^ cell-derived endothelial repair in AVFs via AKT-FoxO1 axis. AKT activation enhances endothelial maturation, thereby attenuating neointimal hyperplasia in AVFs.

**Novelty and Significance:** *What Is Known?:*

- In arteriovenous fistulas (AVFs), abnormal shear stress induces endothelial cell injury, and the resulting neointimal hyperplasia is a major cause of anastomotic stenosis.
- CD34 cells actively participate in vascular endothelial repair.
- PIEZO1 is a mechanoreceptor mediating endothelial sensing of hemodynamic shear stress, contributing to the maintenance of atheroprotective endothelial phenotype under laminar shear stress, whereas its activation induces pro-inflammatory effects under disturbed shear stress.

*What New Information Does This Article Contribute?:*

- CD34 cells participate in repairing endothelial injury induced by abnormal shear stress in AVFs. PIEZO1 knockout in CD34^+^ cells improve endothelial repair and attenuates neointimal hyperplasia in AVF.
- Laminar shear stress induces CD34 downregulation and upregulates VE-cadherin and claudin-5 expression in CD34-high human umbilical vein endothelial cells, whereas oscillatory shear stress upregulates CD34 expression and suppresses VE-cadherin and claudin-5 expression.
- Mechano-stimuli lead to PIEZO1-VEGFR2 complex formation regulating CD34 cell-mediated endothelial repair through the downstream AKT-FoxO1 axis. Abnormal hemodynamic shear stress-induced endothelial injury initiates neointimal hyperplasia in AVFs. The present study identifies PIEZO1 as a key mechanosensor that regulates CD34^+^ cell-derived endothelial repair in response to distinct blood flow patterns. PIEZO1 promotes CD34^+^ cell differentiation into mature ECs for endothelial repair under laminar shear stress, whereas it disrupts the differentiation of CD34^+^ cells into mature endothelium under oscillatory shear stress. Mechanistically, a novel shear stress-sensing complex comprising PIEZO1 and VEGFR2 was identified in regulating flow-induced differentiation of CD34^+^ cells into mature ECs via the AKT-FoxO1 signaling axis, thereby controlling the expression of endothelial maturation markers VE-Cadherin and Claudin-5. These findings define a novel PIEZO1–VEGFR2 mechanotransduction axis in CD34^+^ cell-mediated endothelial repair and support AKT pathway activation as a potential therapeutic strategy against neointimal hyperplasia in AVFs.

## INTRODUCTION

Arteriovenous fistula (AVF) is a clinically preferred long-term hemodialysis access for patients with end-stage renal disease. However, its long-term patency rate is limited by anastomotic stenosis.^1–3^ Hemodynamic flow-related intimal hyperplasia within the arteriovenous fistula is the primary cause of stenosis, maturation failure (33%–62% at 6 months), or poor long-term patency (60%–63% at 2 years).^2^ Up to 50% of AVF patients require percutaneous transluminal angioplasty to alleviate stenosis and associated complications.^4^ Vascular endothelial cell (EC) injury is the key initiating event in the development and progression of anastomotic stenosis.^5,6^ Previous studies have shown that impaired endothelial repair can lead to platelet activation, inflammatory cell recruitment, and smooth muscle cells phenotypic switching, ultimately resulting in neointimal hyperplasia and luminal narrowing.^7,8^ Therefore, improving endothelial repair after arteriovenous anastomosis is essential for preventing anastomotic stenosis.

Endothelial progenitor cells are widely recognized as an important cellular source for endothelial repair. Among them, CD34^+^ cells, a subset of endothelial progenitor cells possessing multilineage differentiation potential, have attracted extensive attention.^9–12^ Previous studies have shown that transplantation of CD34^+^ endothelial progenitor cells enhances endothelial repair, reduces neointimal hyperplasia in artery injury model ^13,14^ and inhibit smooth muscle cell proliferation.^15^ Recent lineage tracing and single-cell RNA sequencing studies have further refined this concept by showing that nonbone marrow-derived CD34^+^ cells are the primary source for endothelial repair.^10^ Despite this progress, the mechanisms governing CD34^+^ cell differentiation remain incompletely understood. Although prior research focusing on the role of biochemical regulators, including inflammatory cytokines, growth factors and metabolic signals has revealed several attractive targets for therapeutic intervention, the incomplete endothelial repair in many vascular diseases indicates additional regulatory mechanisms are involved in CD34^+^ cell fate control. One such likely mechanism is hemodynamic shear stress generated by blood flow, a fundamental biomechanical cue for endothelial homeostasis and vascular remodeling. Given that CD34^+^ cell-participated endothelial repair in large arteries interact with dynamic flow environment, it is rational to hypothesize that shear stress may play a determinant role in regulating CD34^+^ cell fate. This hypothesis is particularly relevant AVFs where vascular remodeling is typically dominated by abnormal shear stress. However, it remains unclear how sustained changes in shear stress are sensed by CD34^+^ cells and transduced into mysterious molecular events to control cell fate and function. Therefore, defining and elucidating the role of shear stress in regulating the differentiation of CD34^+^ cells to mature vascular endothelium is essential for understanding the molecular and cellular basis of impaired endothelial repair of AVF remodeling.

PIEZO1, a highly conserved mechanosensitive ion channel, is a pivotal mechanosensor to transduce the effect of hemodynamic shear stress on EC phenotypes and functions.^16–18^ In mature ECs, PIEZO1 serves as a critical context-dependent regulator of barrier function and vascular integrity. For example, in pulmonary microvessels, pressure-activated PIEZO1 modulates endothelial barrier function by regulating VE-Cadherin phosphorylation and the calcium-dependent protease calpain.^19,20^ However, its role in shear stress-induced differentiation and maturation of CD34^+^ cell-derived ECs remains unexplored. Mature ECs are characterized by the abundant expression of adherens and tight junctional proteins such as VE-cadherin and claudin-5^21^. Notably, VE-cadherin forms a mechanosensory complex with CD31 and vascular endothelial growth receptor-2 (VEGFR2).^22^ As a tyrosine kinase, VEGFR2 integrates signals from hemodynamic shear stress and growth factors to its downstream effectors mainly through PI3K-Akt and MAPK-ERK1/2 pathways.^23–25^ These functions position VEGFR2 as a central signaling hub where biophysical and biochemical cues converge. However, whether VEGFR2 interplays with PIEZO1-mediated mechanotransduction during the differentiation and maturation of CD34^+^ cells remains largely unknown.

In this study, we observed that a large number of CD34^+^ cells was recruited to the injured intima in AVF and participated in endothelial repair. However, the disorganized distribution of ECs and neointimal hyperplasia suggested that the intimal injury was not fully repaired. Based on this observation, we propose that CD34^+^ cells become trapped in a repeated vicious cycle of injury-repair induced by flow disturbance, and that blocking the effect of oscillatory shear stress generated by disturbed flow on CD34^+^ cell differentiation is key to addressing AVF failure. Our in vivo experiments revealed that PIEZO1 knockout in CD34^+^ cells significantly attenuated neointimal hyperplasia in AVFs. The mechanistic study demonstrates a critical role of a shear stress-sensing complex comprising PIEZO1 and VEGFR2 in regulating flow-induced differentiation of CD34^+^ cells into mature ECs via the AKT-FoxO1 signaling axis.

## METHODS

### Data Availability

All supporting data are available within the article and its Data Supplement. For details on the experimental procedures, see the Materials and Methods section in the Data Supplement.

## Result

### CD34^+^ Cells Are Involved in the Endothelial Repair after Arteriovenous Fistula Surgery

Abnormal hemodynamic shear stress is one of the key pathological factors leading to stenosis and arteriovenous fistulas (AVF) failure in the later stages of clinical surgery. To simulate the clinical end-to-side anastomosis of the cephalic vein and radial artery in AVF surgery, we successfully established a mouse AVF model by anastomosing the proximal end of the external jugular vein to the side wall of common carotid artery (Figure 1A). We observed that 4 weeks after AVF establishment, significant adventitial fibrosis and intimal hyperplasia were present at the anastomotic site indicated by HE staining, which are highly similar to the pathological changes observed in human AVF surgery (Figure S1A, S1B). Immunofluorescence *en face* staining indicated that in the normal external jugular vein where blood flow is mostly laminar, only a small number of Cd34-derived (tdTomato^+^) endothelial cells (ECs) were present in the endothelium, and these ECs were regularly aligned along the direction of blood flow. In contrast, at the AVF site, a large number of Cd34-derived ECs were observed, with disorganized alignment and significantly increased intercellular connection distance between endothelial cells (Figure 1B, 1C&1D). This reveals that in AVF model, Cd34^+^ cells participate in repairing injured endothelium caused by disturbed flow. Combining the AVF model with bone marrow transplantation (Figure S1C), we observed consistent results reported by previous study^10^ which showed that nonbone marrow CD34^+^ cells play a crucial role in vascular repair. In Cd34-CreERT2; R26-tdTomato mice receiving C57BL/6J bone marrow, CD34^+^ cells contributed to endothelial repair. In contrast, no tdTomato^+^ cells were detected in C57BL/6J mice receiving bone marrow from Cd34-CreERT2; R26-tdTomato mice (Figure S1D, S1E). To evaluate endothelial barrier function, we detected the fluorescence intensity of VE-Cadherin and Claudin-5 in the AVF model, which was found significantly reduced (Figure 1E,1F & Figure S1F).

**Figure 1.**
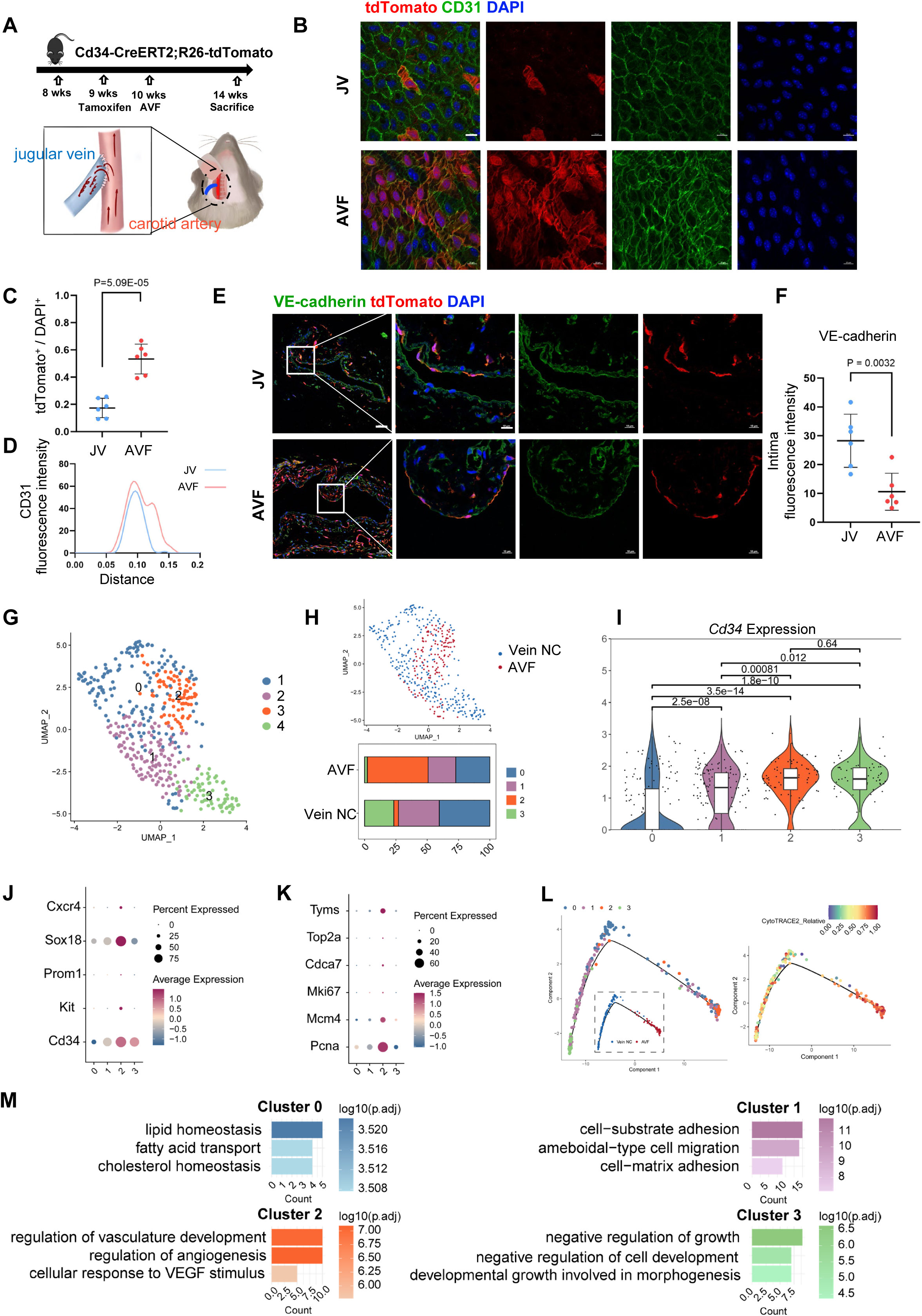
CD34^+^ cells are involved in the endothelial repair after arteriovenous fistula (AVF) surgery. **(A)** Schematic diagram illustrating the lineage tracing strategy and AVF construction in mice. **(B)** En face immunofluorescence staining for tdTomato (red) and CD31 (green) in AVF and the contralateral jugular vein (JV) of Cd34-CreERT2; R26-tdTomato mice. **(C)** Comparisons of the ratio of tdTomato^+^ cells in AVF and contralateral jugular vein. **(D)** Distance of cell junction in AVF and contralateral JV. **(E)** Immunofluorescence staining images of VE-Cadherin (green) and tdTomato (red) in AVF and contralateral JV (overall view scale bar: 50 μm; magnified view scale bar: 10 μm.) **(F)** Summarized data showing intima fluorescence intensity of VE-Cadherin per vessels. **(G)** Focused analysis identifying 4 distinct clusters of endothelial cells (ECs) in normal vein and AVF shown in uniform manifold approximation and projection (UMAP) plot. **(H)** UMAP plot and corresponding bar chart showing the proportion of EC subclusters in normal vein and AVF. **(I)** Violin plot showing Cd34 expression level in 4 EC subclusters. **(J, K)** Dot plots respectively showing the expression levels of proliferation and stemness-associated genes. **(L)** Pseudo-time trajectory of 4 EC subclusters with DDRTree algorithm for dimension reduction, along with the cytotrace2 differentiation score. **(M)** Bar plot showing top enriched gene oncology functions in each EC subcluster. Statistical comparisons were performed using the two-tailed unpaired t-test (C, D, F) and the Mann-Whitney test (I).

To further characterize the functional phenotype of CD34^+^ cells involved in endothelial repair after AVF, we performed single-cell RNA sequencing analysis of cells from both AVF tissues and normal veins. Using UMAP analysis, the cells were classified into nine distinct clusters: fibroblasts (FB, *Col1a1, Col1a2*, etc.), macrophages (*Lyz2, C1qa,* etc.), T cells (*Cd3d, Il7r*, etc.), smooth muscle cells (SMC, *Acta2, Myl9*, etc.), endothelial cells (EC, *Pecam1, Vwf*, etc.), B cells (*Cd79a, Cd37*, etc.), lymphatic endothelial cells (LEC, *Ccl21a, Lyve1*, etc.), plasma cells (*Jchain, Ighm*, etc.), and mesothelial cells (*Mpz*) (Figure S1G,S1H). Among these, Cd34 expression was predominantly high in FB and ECs. Notably, after AVF establishment, Cd34 expression was decreased in FB but increased in ECs (Figure S1I).

To investigate the phenotypic changes in the *Cd34*^high^ EC population before and after arteriovenous fistula surgery, we extracted the EC cluster for secondary dimensionality reduction and subdivided it into 4 subclusters (Figure 1G). We found that both cluster 2 and cluster 3 cells show high *Cd34* expression; however, cluster 2 was enriched in AVF samples, while cluster 3 was predominantly derived from normal control vein samples (Figure 1H,1I). This clustering result suggests that *Cd34*^high^ ECs exhibit distinct gene expression profiles under disturbed flow in AVF compared to those cells under laminar flow in normal veins. To elucidate these phenotypic changes, we examined the expression of endothelial progenitor cell markers, including *Cd34, Kit, Prom1, Sox18*, and *Cxcr4*. We found that cluster 2 cells, exposed to disturbed flow, highly expressed all these markers (Figure 1J). Additionally, cluster 2 cells showed high expression of proliferation markers such as *Pcna, Mcm4, Mki67, Cdca7, Top2a*, and *Tyms*, and exhibited a higher cell cycle score (Figure 1K and Figure S1H), indicating enhanced proliferative capacity. Pseudotime analysis of endothelial subclusters using Monocle2 and CytoTRACE2 revealed that cluster 2 cells were positioned at an early, undifferentiated stage (Figure 1L). Differential gene expression and functional enrichment analysis showed that cluster 0 and 1 cells represent functionally mature endothelial populations, enriched in lipid transport functions and adhesion and migration functions, respectively. In contrast, the two *Cd34*-high clusters (cluster 2 and cluster 3) exhibited distinct functional enrichments. Cluster 2 was enriched in vascular development and angiogenesis, whereas cluster 3 was associated with growth inhibition (Figure 1M).

Based on the above findings, we conclude that shear stress plays a crucial role in regulation of intimal repair mediated by Cd34^+^ cells. Under disturbed flow conditions, Cd34^+^ cells are extensively recruited to repair injured intima. However, persistent disturbed flow environment maintains CD34^+^ cells in a proliferative phenotype, preventing them to become functionally mature endothelial cells with intact barrier function.

### CD34^high^ and CD34^low^ Umbilical Vein Endothelial Cells Exhibit Distinct Gene Expression Phenotypes

To further explore the mechanism by which shear stress regulates Cd34^+^ cell-mediated endothelial repair, we isolated human umbilical vein endothelial cells (HUVECs), and separated the CD34^high^ expressing cell population (defined as CD34^high^ HUVECs) and CD34^low^ HUVECs using flow cytometry (Figure S2A). To verify whether the sorted CD34^high^ HUVECs could represent the early endothelial progenitor cells involved in Cd34^+^ cell-mediated endothelial repair, we compared the gene expression levels of relevant markers in a bulk-sequencing dataset (Figure S2B). Endothelial progenitor cell markers, including *CD34, KIT,* and *KDR*, were highly expressed in CD34^high^ HUVECs, whereas endothelial cell maturation markers, such as the cadherin gene *CDH5*, tight junction gene *CLDN5*, integrin genes *ITGB1, ITGB3*, and *ITGA5*, and endothelial nitric oxide synthase *NOS3*, were expressed at low level in CD34^high^ HUVECs (Figure S2C). These data indicate that the separated CD34^high^ HUVECs exhibit features of early endothelial progenitor cells and can serve as a model for studying CD34^+^ cell-mediated endothelial repair.

Next, we compared gene expression of mechanosensitive transcription factors and mechanosensors between CD34^high^ and CD34^low^ HUVECs. CD34^high^ HUVECs predominantly express high level of developmental transcription factors, such as *POU2F2* and *TCF15*; as well as osteogenesis-related transcription factors, including *ATF3, RUNX1, SMAD1*, and *SMAD5*. In contrast, CD34^low^ HUVECs showed higher expression of shear stress-responsive transcription factors, such as *KLF2, KLF4,* and *MEF2C*, which are known to in maintain endothelial barrier function under laminar shear stress (Figure S2D). Consistently, mechanosensor genes, including *DDR1, CD44, PIEZO1, PLXD1*, and *HEG1*, were also more highly expressed in CD34^low^ HUVECs (Figure S2E). We stimulated CD34^low^ and CD34^high^ HUVECs with laminar shear stress (LSS) and oscillatory shear stress (OSS) respectively, and found that KLF2 was more strongly upregulated in CD34^low^ HUVECs under LSS (Figure S2F) and the proinflammatory adhesion molecule VCAM-1 was also more strongly upregulated in CD34^low^ HUVECs under OSS (Figure S2G). These results suggest that CD34^high^ HUVECs exhibit distinct mechanosensitive profile and are likely to respond differently to biomechanical cues compared to mature endothelial cells characterized in previous studies.

### Shear Stress Regulates the Maturation of CD34^high^ HUVECs

We seeded CD34^high^ HUVECs to flow chamber, and exposed them to a distinct shear stress stimulation to simulate the in vivo hemodynamic environment (Figure 2A). After 4-hour exposure to LSS, we detected an increased mRNA expression of *KLF2* and *HMOX1*, known flow-induced genes, in CD34^high^ HUVECs (Figure S2H), suggesting successfully flow stimulation. We observed a significant decrease in the mRNA expression of the endothelial progenitor cell markers *CD34* and *KIT*, while significantly increased mRNA expression of mature EC markers *CDH5*, *VWF*, *FLT1* and *CLDN5* (Figure 2B). Following 24-hour LSS stimulation, CD34 protein expression was markedly reduced, and VE-Cadherin and Claudin-5 expression was significantly elevated (Figure 2C). Additionally, KLF2 expression, a positive control for stimulation, was upregulated (Figure 2C). Immunofluorescence analysis confirmed that CD34 fluorescence intensity significantly decreased after 24-hour LSS stimulation, and the proportion of KI67^+^ cells was significantly reduced (Figure 2D, 2E). These results indicate that LSS promotes the differentiation, maturation, and growth arrest of CD34^high^ HUVECs.

**Figure 2:**
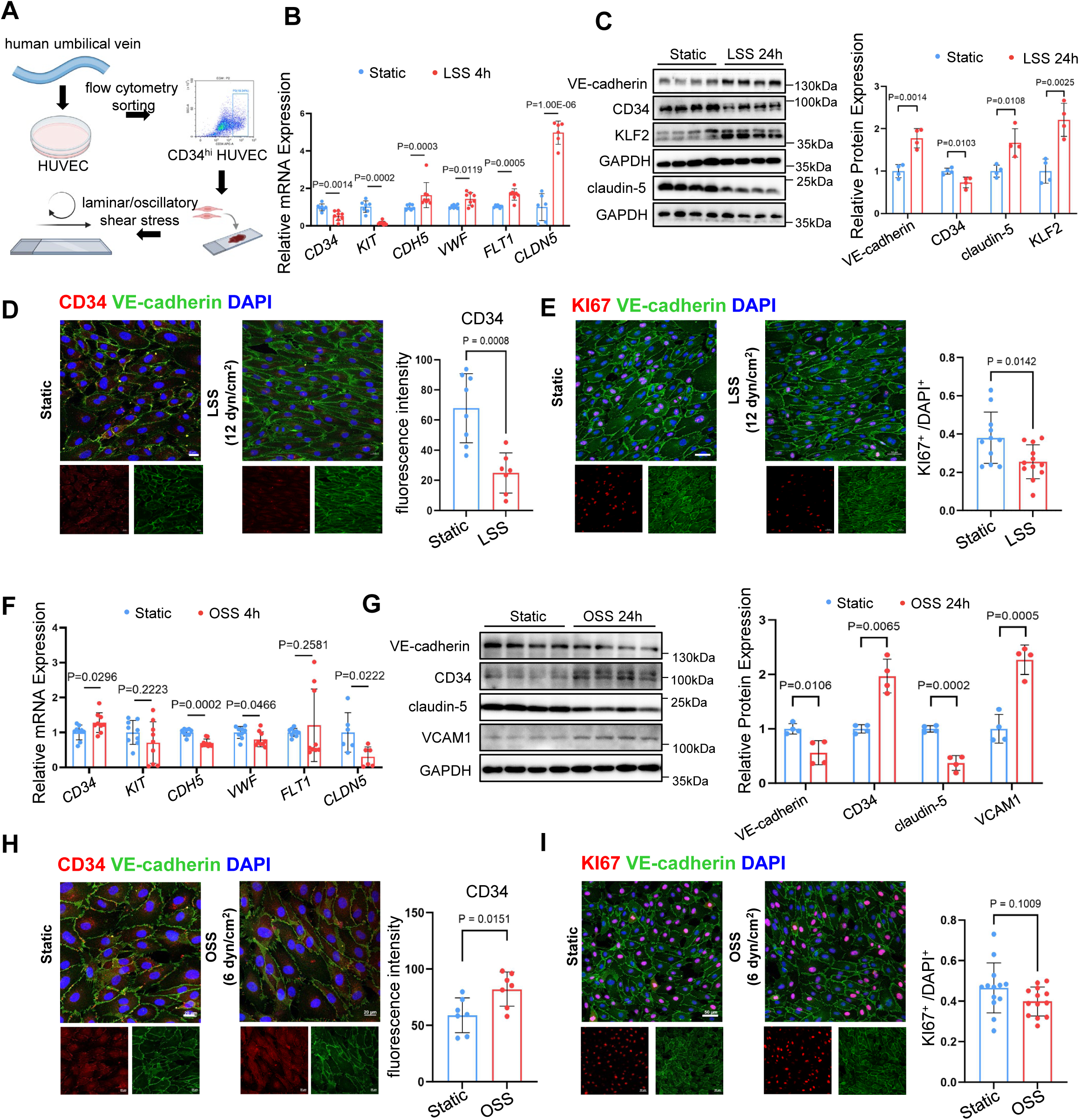
shear stress regulates maturation of CD34^high^ human umbilical vein endothelial cells (HUVECs). **(A)** Schematic diagram illustrating the workflow of in vitro cell sorting and shear stress stimulation. **(B)** qPCR analysis with quantification normalized to GAPDH in CD34^high^ HUVEC exposed to laminar shear stress (LSS, 12 dynes/cm^2^) for 4 hours or static control. **(C-E, G-I)** CD34^high^ HUVECs were exposed to LSS at 12 dyn/cm^2^ **(C-E)** or oscillatory shear stress (OSS, 6 dyn/cm^2^) (G-I) for 24 hours and static control. **(C, G)** Western blotting analysis with quantification normalized to GAPDH. **(D, H)** CD34^high^ HUVECs were stained for CD34 (red), VE-Cadherin (green), and DAPI. Bar graph showing summarized data of CD34 fluorescence intensity. Scale bar: 20 μm. **(E, I)** CD34^high^ HUVECs were stained for KI67 (red), VE-Cadherin (green), and DAPI. Bar graph showing the ratio of KI67 to DAPI. Scale bar: 50 μm. **(F)** qPCR analysis with quantification normalized to GAPDH in CD34^high^ HUVEC exposed to OSS at 6 dynes/cm^2^ for 4 hours or static control. Statistical comparisons were performed using the two-tailed unpaired t-test (CD34 and CLDN5 in panel B; panels C, D, and H; CD34, KIT, VWF, and CLDN5 in panel F; VE-Cadherin, Claudin-5, and VCAM1 in panel G), the two-tailed unpaired Welch’s t-test (KIT, VWF, and FLT1 in panel B; CD34 in panel G), and the Mann-Whitney test (CDH5 in panel B; VWF and FLT1 in panel F).

On the other hand, compared with the static group, OSS stimulation slightly increased *CD34* mRNA expression in CD34^high^ HUVECs, while no difference was observed in *KIT* expression. The mRNA levels of the EC maturation markers *CDH5*, *VWF* and *CLDN5* were decreased (Figure 2F). As positive control, the mRNA expression of *KLF2* was decreased and *VCAM1* was increased (Figure S2I). After 24-hour OSS stimulation, CD34 protein expression was increased compared to the control group, whereas VE-Cadherin and Claudin-5 protein expression levels were reduced (Figure 2G). Upregulated VCAM-1 expression was detected to show successful induction of OSS (Figure 2G). Immunofluorescence analysis confirmed that CD34^high^ HUVECs maintained higher CD34 expression after 24-hour exposure to OSS (Figure 2H). There was no significant difference in the proportion of KI67^+^ cells between OSS and static groups (Figure 2I), indicating that CD34^high^ HUVECs under OSS exhibited a proliferative capacity similar to that under static culture conditions. EdU staining similarly demonstrated that LSS inhibited whereas OSS sustained proliferation (Figure S3A). Together, these findings indicate that OSS maintains CD34^high^ HUVECs at an incompletely differentiated state.

### PIEZO1 Mediates the Effect of Shear Stress on the Maturation of CD34^high^ HUVECs

Mechanosensors serve as a bridge connecting extracellular biomechanical signals to intracellular biological responses^26–28^. We compared the expression of mechanosensors in EC subclusters from single-cell RNA sequencing data and found that cells with higher PIEZO1 expression tend to show higher CD34 expression (Figure S3B) while others mechanosensitive proteins showed no clear correlation with CD34 expression. Furthermore, treatment with the PIEZO1 inhibitor GsMTx4 upregulated CD34 expression in CD34^high^ HUVECs under LSS stimulation, although it had no significant effect on CD34 expression under OSS conditions (Figure S3C).

To explore the potential role of PIEZO1 in the LSS-induced maturation of CD34^high^ HUVECs, we transduced HUVECs with lentivirus carrying shRNA targeting PIEZO1 and validated a more than 60% knockdown at mRNA and protein level (Figure 3A, 3B). CD34^high^ HUVECs with PIEZO1 knockdown and Control cells transduced with scrambled shRNA (control) were subjected to LSS for 24 hours. Western blotting results showed that in PIEZO1 knockdown group, the inhibitory effects of LSS on CD34 expression were attenuated, while VE-cadherin and claudin-5 showed decreased expression compared to scramble group (Figure 3C, 3D). Under LSS conditions, PIEZO1 knockdown increased CD34 fluorescence intensity and the proportion of Ki67-positive cells compared to control group (Figure 3E-3G), indicating that PIEZO1 mediates the LSS-induced differentiation and maturation of CD34^high^ HUVECs.

**Figure 3:**
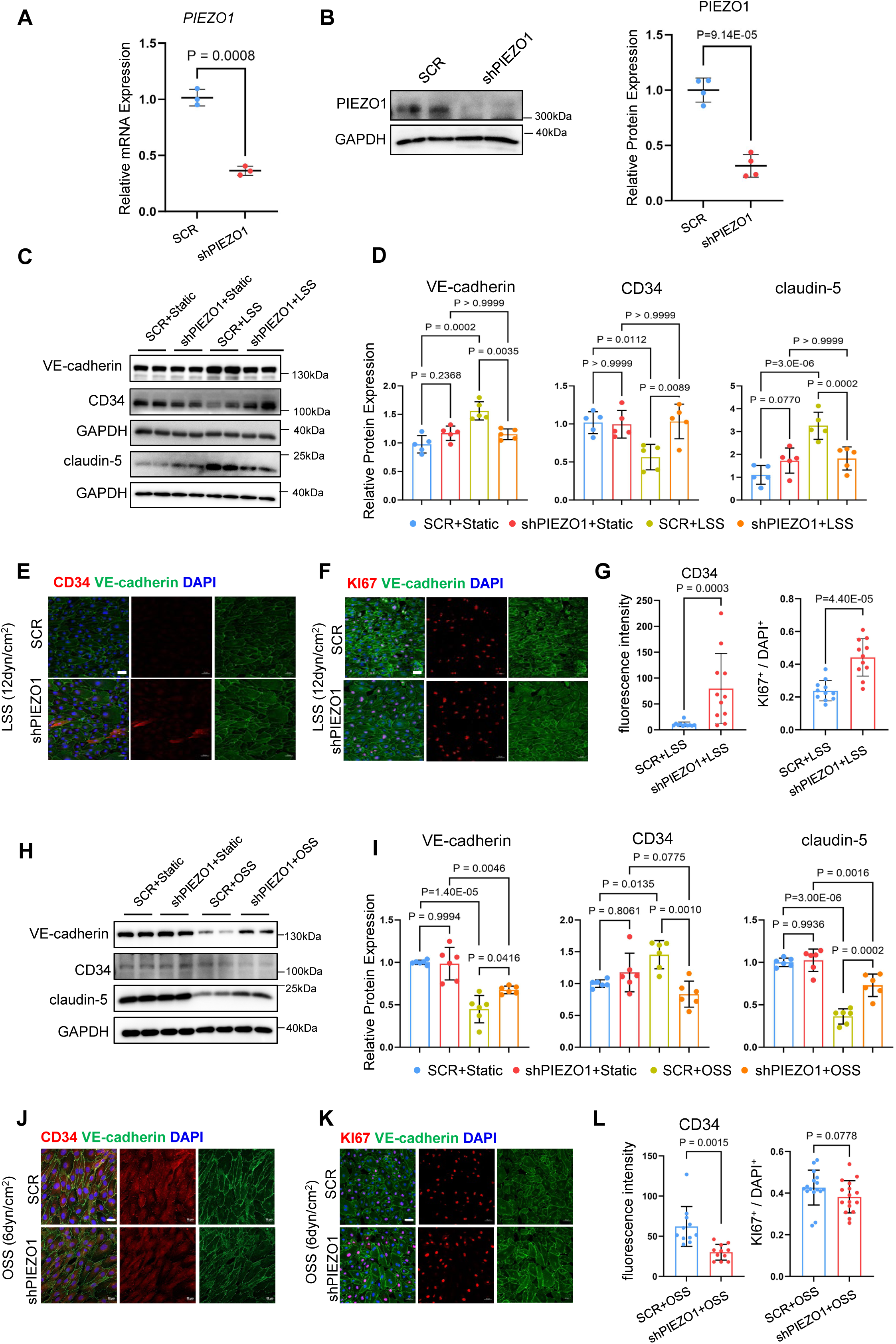
PIEZO1 mediates shear stress-induced effect on maturation of CD34^high^ HUVECs. **(A-B)** LV-shPIEZO1 knock down PIEZO1 mRNA (A) and protein (B) levels in CD34^high^ HUVECs. **(C-L)** CD34^high^ HUVECs transduced with lentivirus carrying LV-shPIEZO1 or LV-scramble (SCR) for 72h followed by exposure to laminar shear stress (LSS, 12 dyn/cm^2^) **(C-G)** or oscillatory shear stress (OSS, 6 dyn/cm^2^) **(H-L)** for 24 hours and static control. **(C-D, H-I)** Western blotting analysis with quantification normalized to GAPDH. **(E, J)** CD34^high^ HUVECs were stained for CD34, VE-Cadherin, and DAPI. **(G, L left panel)** Bar graph showing summarized data of CD34 fluorescence intensity. Scale bar: 20 μm. **(F, K)** CD34^high^ HUVECs were stained for KI67, VE-Cadherin, and DAPI. **(G, L right panel)** Bar graph showing the ratio of KI67 to DAPI. Scale bar: 50 μm. Statistical comparisons were performed using the two-tailed unpaired t-test (B; KI67/DAPI in panel G), the two-tailed unpaired Welch’s t-test (A; CD34 in panel L), the Mann-Whitney test (KI67/DAPI in panel L; CD34 in panel G), and two-way ANOVA with Bonferroni correction (D, I).

To investigate the role of PIEZO1 in maintaining incompletely differentiated state in CD34^high^ HUVECs under disturbed flow conditions, we exposed the cells to OSS for 24 hours and found attenuated maintenance of CD34 expression, and diminished inhibitory effects on VE-Cadherin and Claudin-5 expression by OSS in PIEZO1 knockdown group (Figure 3H, 3I). Under OSS conditions, CD34 fluorescence intensity was significantly lower in PIEZO1 knockdown group compared to control (Figure 3J, 3L), while no difference was observed in the proportion of Ki67^+^ cells between groups (Figure 3K, 3J). These findings suggest that PIEZO1 plays distinct roles under OSS and LSS, contributing to the maintenance of stemness under OSS without affecting proliferative capacity of in CD34^high^ HUVECs.

### *Piezo1* Knockout in CD34^+^ Cells Promotes Endothelial Repair and Attenuates Intimal Hyperplasia

To investigate the functional role of PIEZO1 in CD34^+^ cell-mediated endothelial repair in AVF, we generated Cd34-CreERT2; R26-tdTomato; Piezo1^flox/flox^ mice (CKO), enabling conditional knockout of *Piezo1* in CD34-expressing cells (Figure 4A and Figure S3D). Single-cell RNA sequencing of AVF tissue revealed that *Piezo1* is predominantly expressed in ECs and lymphatic ECs (Figure 4B). Immunofluorescence staining showed that tdTomato^+^ and PIEZO1^+^ cells primarily co-expressed CD31, indicating that although CD34^+^ cells have the potential to differentiate into fibroblasts and immune cells, they mainly contribute to endothelial repair by differentiating into ECs in AVF model (Figure 4C).

**Figure 4:**
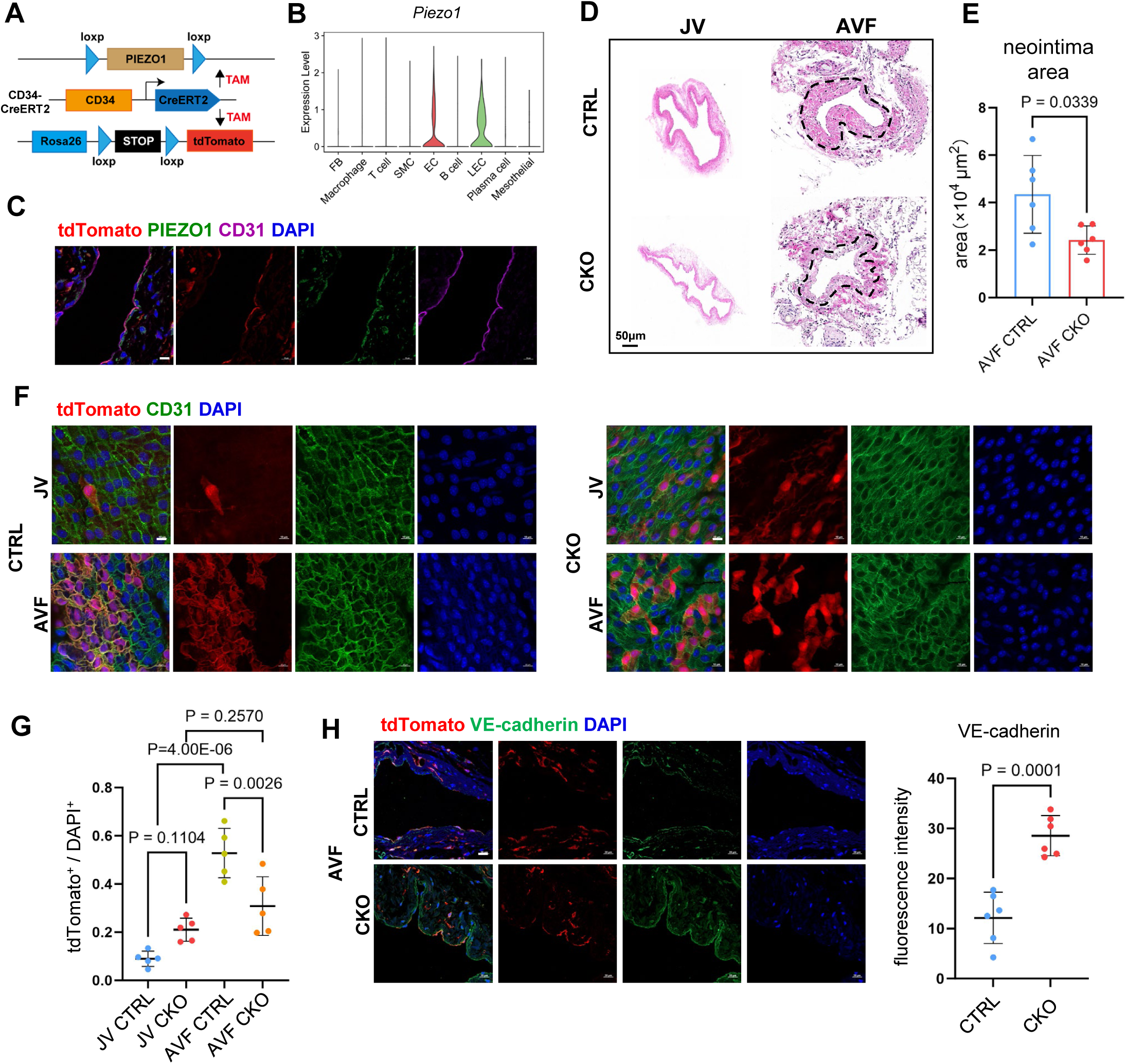
PIEZO1 conditional knockout in CD34^+^ cells improves endothelial repair and attenuates neointimal hyperplasia of AVF. (A) Schematic diagram illustrating the construction strategy of CD34-CreERT2; R26-tdTomato; Piezo1^flox/flox^ mice. (B) Violin plot showing Piezo1 expression level in each cell cluster from AVF and normal vein. (C) Immunofluorescence images for tdTomato (red), PIEZO1 (green) and CD31 (purple) in AVF, Scale bar: 10 μm. (D-E) H&E staining images of AVF and jugular vein (JV) from Cd34-CreERT2; R26-tdTomato (CTRL) and CD34-CreERT2; R26-tdTomato; Piezo1^flox/flox^ (CKO) mice, with quantifications of neointima areas. Scale bar: 50 μm. (F) *En face* immunofluorescence images for tdTomato (red) and CD31 (green) in AVF and contralateral JV from CTRL and CKO mice. Scale bar: 10 μm. (G) Dot plot graph showing the ratio of tdTomato^+^ cell to DAPI. (H) Immunofluorescence images for tdTomato (red), VE-Cadherin (green) and DAPI in AVF from CTRL and CKO mice, Scale bar: 20 μm. Statistical comparisons were performed using the two-tailed unpaired Welch’s t-test (E), two-way ANOVA with Bonferroni correction (G), and two-tailed unpaired t-test (H).

In AVF model, *Piezo1*-CKO mice exhibited significantly reduced neointimal area compared to control mice (Figure 4D), suggesting that disturbed flow-induced neointimal formation was attenuated following Cd34^+^ cell-specific Piezo1 knockout. To further assess changes in endothelial repair, *en face* staining of the fistulas was performed. The results showed that ECs in *Piezo1*-CKO mice exhibited a more spindle-shaped morphology compared to those in controls, and the number of tdTomato^+^ cells was significantly reduced (Figure 4F, 4G). This indicates that Cd34^+^ cells differentiated into more mature ECs, forming a relatively intact endothelial barrier, thereby mitigating the effects of disturbed flow on neointimal hyperplasia.

Although our in vitro experiments showed that PIEZO1 knockdown enhanced the proliferative capacity of CD34^high^ HUVECs under LSS, we did not observe significant difference in tdTomato^+^ cell number between CKO and control mice in normal veins in vivo, likely due to insufficient recruitment signals from intact endothelium in normal veins. After Piezo1 deletion, ECs showed higher levels of VE-Cadherin and Claudin-5 indicated by immunofluorescence staining (Figure 4H and Figure S3E), consistent with above mentioned role of PIEZO1 in mediation of disturbed flow-induced disruption of junctional maturation.

### The PI3K-AKT Signaling Pathway was Significantly Altered Following PIEZO1 Knockdown

To elucidate how PIEZO1 mediates the effect of shear stress on the differentiation and maturation of CD34^+^ cells, we performed a virtual Piezo1 knockout in EC subclusters from single-cell RNA sequencing data using the scTenifoldKnk package. This approach allowed us to enrich for genes co-expressed with Piezo1 and predict its potential biological functions. Because cluster 2 is primarily from AVF and cluster 3 from normal vein, the virtual Piezo1 knockout recapitulated the biological effects of Piezo1 under disturbed (cluster 2) and laminar (cluster 3) shear stress, respectively.

We found that in cluster 2 (mainly from AVF group), virtual Piezo1 knockout downregulated TNF, NF-κB, and TGF-beta signaling pathways. This is consistent with previous reports that Piezo1 mediates disturbed flow-induced inflammatory phenotypes and upregulation of mesenchymal transition genes in ECs. Piezo1 has also been shown to influence mesenchymal stem cells via activation of the MAPK (MEK/ERK) signaling pathway. In cluster 3 (mainly from vein NC group), virtual Piezo1 knockout resulted in downregulation of MAPK, PI3K-AKT, Ras, and mTOR signaling pathways (Figure 5A). In addition, we performed bulk RNA sequencing on CD34^high^ HUVECs with PIEZO1 knockdown or SCR controls under laminar or disturbed shear stress conditions. PIEZO1 knockdown produced opposing PI3K–AKT responses: AKT signaling increased under laminar shear but decreased under disturbed shear (Figure 5B).

**Figure 5:**
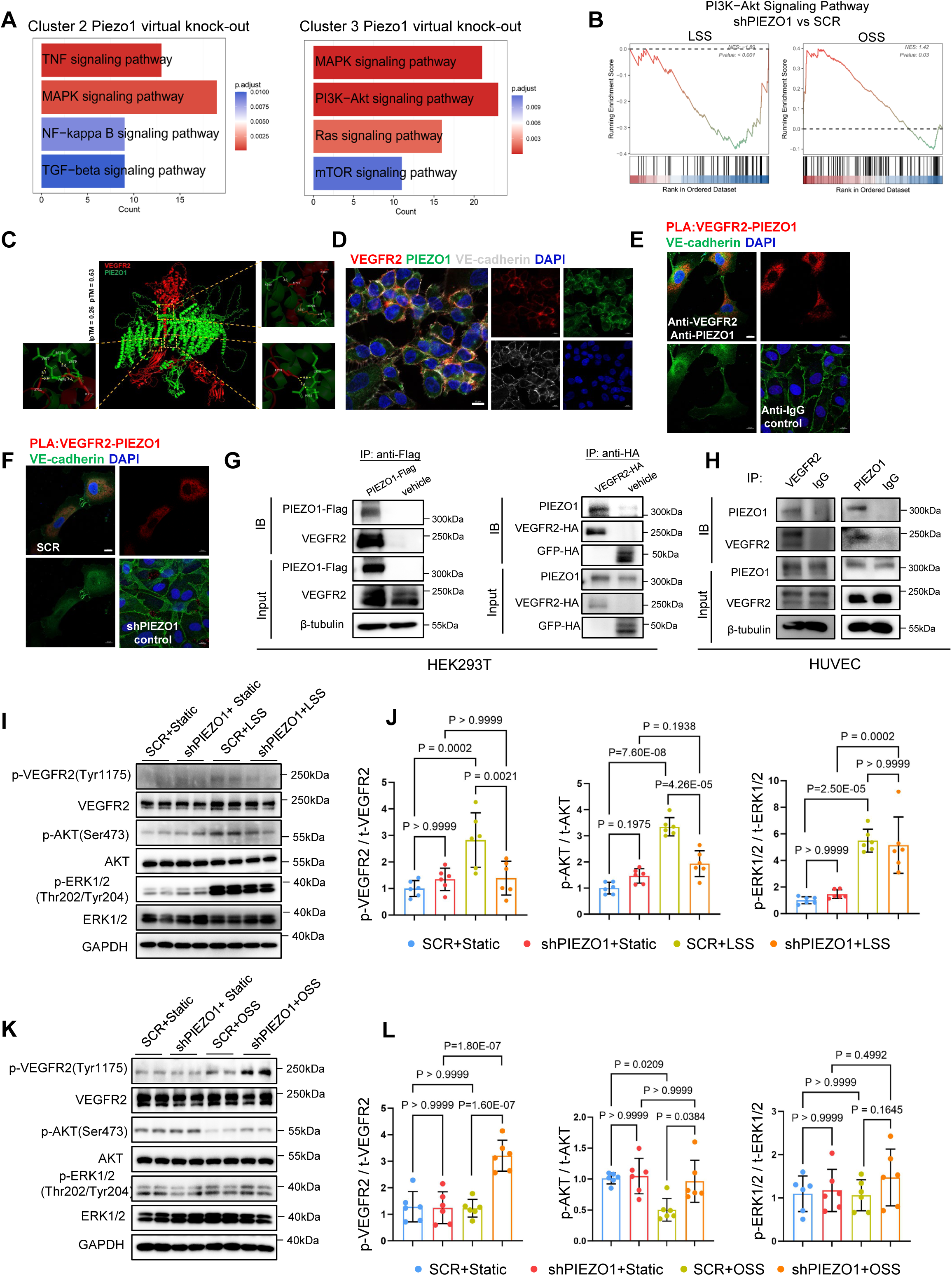
PIEZO1 forms a complex with VEGFR2 to mediate shear stress-induced effect on AKT signaling. **(A)** Bar charts illustrating the functional enrichment of genes regulated by in virtual Piezo1 knockout in EC 2&3 subclusters of Fig 1G. **(B)** Gene set enrichment analysis (GSEA) showing the functional enrichment of CD34^high^ HUVECs treated with LV-shPIEZO1 compared to LV-scramble (SCR) under laminar (LSS, left) and oscillatory (right, OSS) shear stress conditions. **(C)** Structural prediction using AlphaFold3 indicates a potential interaction between VEGFR2 and PIEZO1. **(D)** Immunofluorescence staining images of VEGFR2 (red), PIEZO1 (green) and VE-Cadherin (grey) in CD34^high^ HUVECs. Scale bar: 10μm. **(E-F)** Proximity ligation assay (PLA) of VEGFR2-PIEZO1 probe (red) and immunofluorescence staining images of VE-Cadherin (green). Scale bar: 10μm. **(G-H)** Western blots showing co-immunoprecipitation data obtained from HEK293T cells (G) co-transfected with PIEZO1-Flag and VEGFR2-HA plasmids and HUVEC (H). **(I-L)** Western blots showing phosphorylated and total level of VEGFR2 and its downstream effector proteins (AKT and ERK1/2) in CD34^high^ HUVECs treated with LV-shPIEZO1 or LV-SCR, following exposure to LSS (I-J) or OSS (K-L) for 30 min and static control. Statistical comparisons were performed using two-way ANOVA with Bonferroni correction (J, L).

### PIEZO1 Forms a Complex with VEGFR2 to Mediate the Effect of Shear Stress on AKT Signaling

AKT signaling is a major effector downstream of growth factor signaling pathways. In ECs, activation of vascular endothelial growth factor receptor 2 (VEGFR2) promotes AKT phosphorylation. In addition to its role in growth factor signaling, VEGFR2 acts as a critical mechanosensor in ECs by transducing hemodynamic forces into biochemical signals that regulate vascular permeability and angiogenesis ^29^. VEGFR2 is also an essential component of a specialized junctional mechanosensory complex comprising PECAM-1 and VE-Cadherin ^30^. Based on this prior knowledge and our virtual Piezo1 knockout data, we hypothesized that PIEZO1 likely forms a shear stress-sensing complex with VEGFR2 to control the differentiation and maturation of CD34^+^ cells under different flow patterns. Using AlphaFold3, we predicted the potential interaction between a PIEZO1 monomer and a VEGFR2 monomer at the membrane surface. The prediction suggested a possible spatial interaction, with an ipTM score of 0.26, which is comparable to the predicted interaction score (ipTM=0.31) for the known interacting pair, PIEZO1 and PECAM1^31^ (Figure 5C).

To validate the PIEZO1-VEGFR2 complex, we first observed colocalization of PIEZO1 and VEGFR2 on the membrane surface of CD34^high^ HUVECs by immunofluorescence staining (Figure 5D). Proximity ligation assay (PLA) further revealed the *in situ* protein interaction between PIEZO1 and VEGFR2(Figure 5E). Compared to the control groups, cells incubated with PIEZO1 and VEGFR2 antibodies exhibited distinct PLA punctate signals. Consistently, when PIEZO1 knockdown markedly reduced the PLA punctate signals compared to the SCR control group (Figure 5F). Co-immunoprecipitation in HEK293T cells co-transfected with PIEZO1-Flag and VEGFR2-HA confirmed reciprocal interaction between PIEZO1 and VEGFR2 (Figure 5G). Endogenous co-immunoprecipitation in HUVEC further confirmed PIEZO1 and VEGFR2 form a complex (Figure 5H).

Given that previous studies have shown that shear stress regulates the phosphorylation of VEGFR2 at the Tyrosine 1175 (Try1175) site, we sought to investigate whether the PIEZO1-VEGFR2 complex transduces shear stress signals to VEGFR2 downstream pathways. In PIEZO1 knockdown CD34^high^ HUVECs, we found that the PIEZO1-VEGFR2 complex is critical for mediating LSS-induced VEGFR2 Try1175 phosphorylation and subsequent AKT pathway activation, whereas Erk1/2 signaling was unaffected (Figure 5I, 5J). Under disturbed flow conditions, PIEZO1 knockdown significantly increased VEGFR2 Try1175 phosphorylation and AKT signaling activation, while Erk1/2 signaling remained unchanged (Figure 5K, 5L). These findings indicate that the PIEZO1-VEGFR2 complex plays an inhibitory role in shear stress-mediated VEGFR2 Try1175 phosphorylation and AKT activity.

### PIEZO1 Mediates the Effect of Shear Stress on VEGFR2 Endocytic Signal Transduction Patterns

Phosphorylation of VEGFR2 at the Tyr1175 site is closely associated with endocytic signaling and plays a critical role in regulating gene expression programs related to cell migration, proliferation, and homeostasis^32^. Our bulk RNA sequencing data showed that endocytosis-related functions was significant downregulated in shPIEZO1 group of CD34^high^ HUVECs exposed to LSS compared to the SCR group (Figure S4A). Therefore, we hypothesize that PIEZO1 is likely to regulate VEGFR2 endocytic signaling, thereby modulating shear stress-induced AKT signaling.

We conducted membrane/cytoplasmic fractionation of PIEZO1 knockdown or control CD34^high^ HUVECs exposed to LSS or OSS. LSS application significantly increased the ratio of cytoplasmic to membrane VEGFR2 expression compared to static conditions, whereas no difference of VEGFR2 translocation was observed in shPIEZO1 group (Figure S4B, S4C). Under OSS, shPIEZO1 group showed a significant increase of cytoplasmic-to-membrane VEGFR2 ratio comparing with SCR group (Figure S4D, S4E). We next used an antibody feeding assay to directly visualize changes in VEGFR2 spatial localization. Under LSS, VEGFR2 in SCR group exhibited a punctate cytoplasmic distribution, with significantly reduced colocalization with the membrane protein VE-Cadherin compared to static conditions; in contrast, shPIEZO1 group showed increased colocalization of VEGFR2 with VE-Cadherin compared to the SCR group (Figure S4F, S4G). Under OSS, VEGFR2 distribution in SCR group was similar to that under static conditions, while shPIEZO1 group displayed a punctate cytoplasmic pattern, with significantly increased colocalization with VE-Cadherin compared to static conditions (Figure S4H, S4I). These results corroborate our hypothesis that PIEZO1 plays a crucial role in mediating shear stress-induced effects on VEGFR2 endocytic signaling in CD34^high^ HUVECs.

### The AKT-FoxO1 Axis Mediates Shear Stress-Induced Effect on Maturation of CD34^high^ HUVECs

We next sought to extrapolate our findings obtained from cellular studies to AVF models. We compared the fluorescence intensity of p-AKT in AVF model between Piezo1-CKO and control mice and found that p-AKT signal intensity in the vascular intima was significantly higher in *Piezo1*-CKO mice (Figure 6A), confirming PIEZO1 mediates the inhibitory effect of disturbed flow on AKT signaling. To test whether inhibited AKT signaling is a key factor in the immature endothelial repair mediated by CD34^+^ cells in the AVF model, we first investigated the role AKT signaling in regulating CD34^high^ HUVEC maturation. Under LSS, CD34 fluorescence intensity was significantly increased after treatment with AKT inhibitor MK2206 (Figure 6B). Western blotting results showed that LSS-regulated expression of CD34, VE-cadherin and claudin-5 were mostly reversed by MK2206 treatment (Figure 6C, 6D), suggesting a key role of AKT in mediating flow-induced maturation-linked gene expression program in CD34^high^ HUVECs. Conversely, CD34^high^ HUVECs exposed to OSS upon treatment with AKT activator SC79 showed a significant decrease in CD34 fluorescence intensity (Figure 6E, 6F). OSS-induced effect on CD34, VE-cadherin and claudin-5 expression was abolished by SC79 as confirmed by western blotting data (Figure 6G, 6H). Under static conditions, AKT activation by SC79 reduced CD34 expression but did not induce changes in VE-cadherin or claudin-5 expression (Figure S5A). These results demonstrate that the AKT activity is key to shear stress-mediated maturation of CD34^high^ HUVECs.

**Figure 6:**
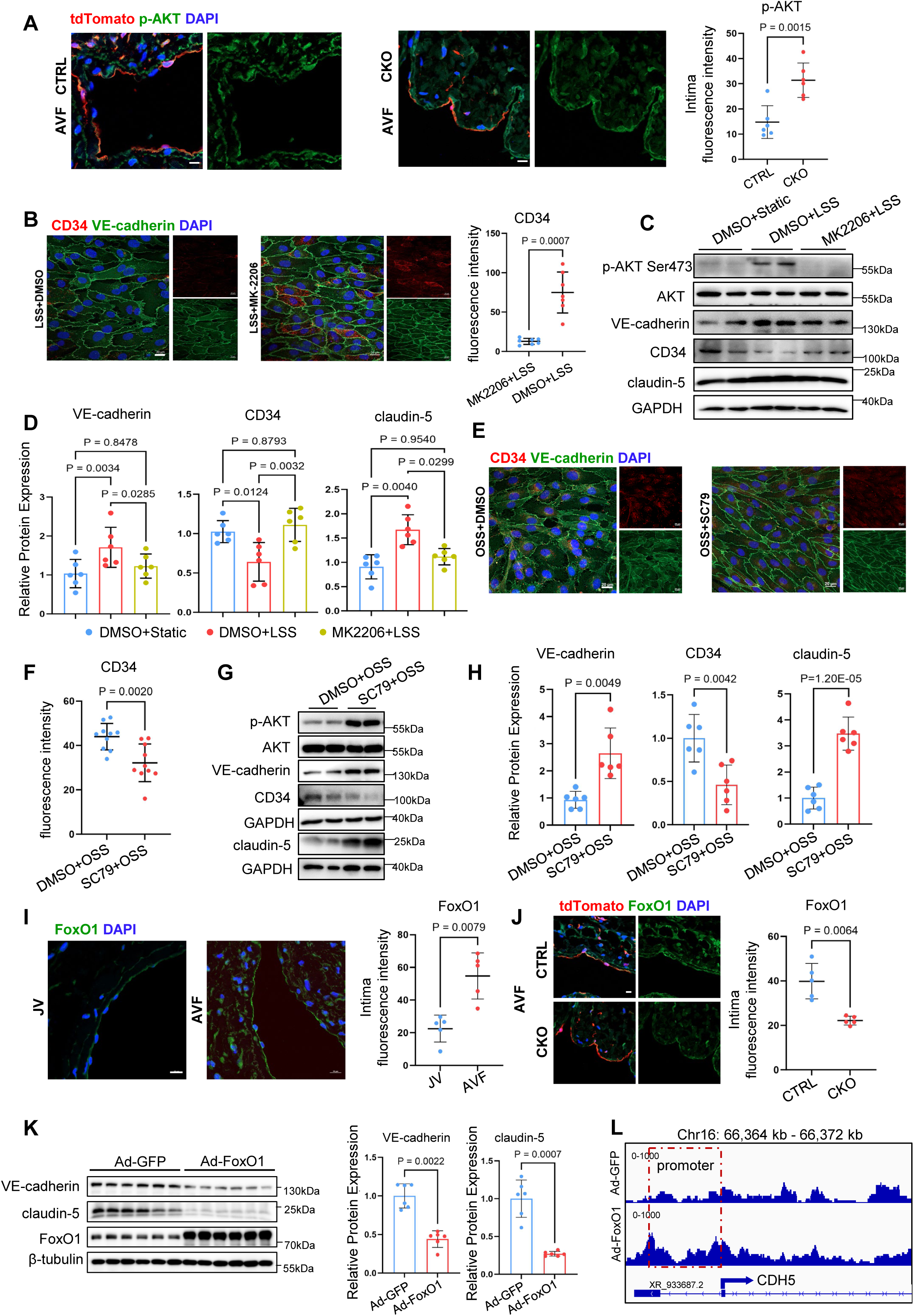
The AKT-FoxO1 axis regulates maturation of CD34^+^-derived cells. **(A)** Immunofluorescence images of p-AKT (green) and tdTomato (red) in AVF, with comparison of p-AKT fluorescence intensity per vessels. Scale bar: 20 μm. **(B-D)** CD34^high^ HUVECs exposed to laminar shear stress (LSS) with MK-2206 or DMSO for 24h. **(E-H)** CD34^high^ HUVECs exposed to oscillatory shear stress (OSS) with SC79 or DMSO for 24h. **(B, E-F)** Immunofluorescence images of CD34^high^ HUVECs for CD34 (red), VE-Cadherin (green) and DAPI. Graph displays CD34 fluorescence intensity. Scale bar: 20 μm. **(C-D, G-H)** Western blot analyses with quantification normalized to GAPDH. **(I)** Immunofluorescence images of FoxO1 (green) in AVF and contralateral jugular vein (JV), with comparison of fluorescence intensity of FoxO1 per vessels. Scale bar: 10 μm. **(J)** Immunofluorescence staining images of FoxO1 (green) and tdTomato in the AVF from CTRL and CKO mice, with comparison of fluorescence intensity of FoxO1 per vessels. Scale bar: 20 μm. **(K)** Western blotting analysis showing the expression of VE-Cadherin and Claudin-5 in CD34^high^ HUVECs transduced with Ad-GFP (green fluorescent protein) and Ad-FoxO1, with quantification normalized to β-tubulin. **(L)** The CUT&Tag peak tracks showing changes in chromatin accessibility at the CDH5 gene locus in HUVECs treated with Ad-GFP and Ad-FoxO1. Statistical comparisons were performed using the two-tailed unpaired t-test (A, F; CD34 and Claudin-5 in panel H), the two-tailed unpaired Welch’s t-test (B; VE-Cadherin in panel H; J; Claudin-5 in panel K), the Mann-Whitney test (I; VE-Cadherin in panel K), and two-way ANOVA with Bonferroni correction (D).

To identify transcriptional regulators downstream of AKT potentially involved in the differentiation of CD34-derived cells into mature ECs, we analyzed differential expression changes of transcription factors between endothelial cluster 2 and cluster 3 using single-cell RNA sequencing data. We found that transcription factors such as *Sox7*, *Sox17*, *Klf6*, and *Foxo1*, which are associated with shear stress and angiogenesis, were upregulated in cluster 2 (Figure S5B). Among these, *Foxo1* has been previously reported to be linked to AKT signaling and involved in a variety of cellular processes including proliferation, differentiation.^33,34^ To investigate whether shear stress-induced AKT signaling also regulates FoxO1 in CD34^high^ HUVECs, we applied short-duration LSS and OSS together with pharmacological inhibition or activation of AKT signaling. Our results showed that LSS promoted FoxO1 phosphorylation, which was inhibited by MK2206 (Figure S5C). In contrast, OSS suppressed FoxO1 phosphorylation, and AKT activation under both static and OSS conditions promoted FoxO1 phosphorylation (Figure S5D). FoxO1 phosphorylation facilitates its cytoplasmic translocation, leading to proteasomal degradation and decreased transcriptional activity^35^, suggesting that FoxO1-mediated transcriptional program may function downstream of AKT signaling during CD34^high^ HUVEC maturation. We observed significantly increased FoxO1 fluorescence intensity in AVF model (Figure 6I), and *Piezo1*-CKO markedly reduced FoxO1 fluorescence intensity in AVF intima compared to control mice (Figure 6J).

To confirm that disturbed flow-induced upregulation of FoxO1 directly inhibits the maturation of CD34^high^ HUVECs, we overexpressed FoxO1 in CD34^high^ HUVECs using Adtrack-CMV-FoxO1. VE-Cadherin and Claudin-5 expression was suppressed by FoxO1 overexpression (Figure 6K). To disclose whether FoxO1 directly regulates VE-Cadherin and Claudin-5 in CD34^high^ HUVECs, we used CUT&Tag analysis and found that the peak signal in the CDH5 promoter region was increased following FoxO1 overexpression, indicating that FoxO1 directly binds to CDH5 promoter to negatively regulates its expression (Figure 6L). In summary, our data demonstrate that PIEZO1-VEGFR2 complex senses shear stress to transduce extracellular biomechanical cues into biological signals to regulate the maturation of CD34^high^ HUVECs through the AKT-FoxO1 axis.

### Human AVF Endothelium Exhibits CD34 Expression and Decreased AKT Signaling

To determine whether the endothelial phenotype in human AVF aligns with what we’ve observed in the mouse model, we analyzed single-cell RNA sequencing data of ECs from human AVF specimens and clinically-matched control veins. Among five endothelial subpopulations (Figure 7A), we identified a distinct subcluster characterized by high expression level of genes associated with endothelial development and angiogenesis, including *CD34*, *KIT*, *MEF2C* and *SOX4*, suggesting an endothelial progenitor-like identity (Figure 7B, 7C).

**Figure 7:**
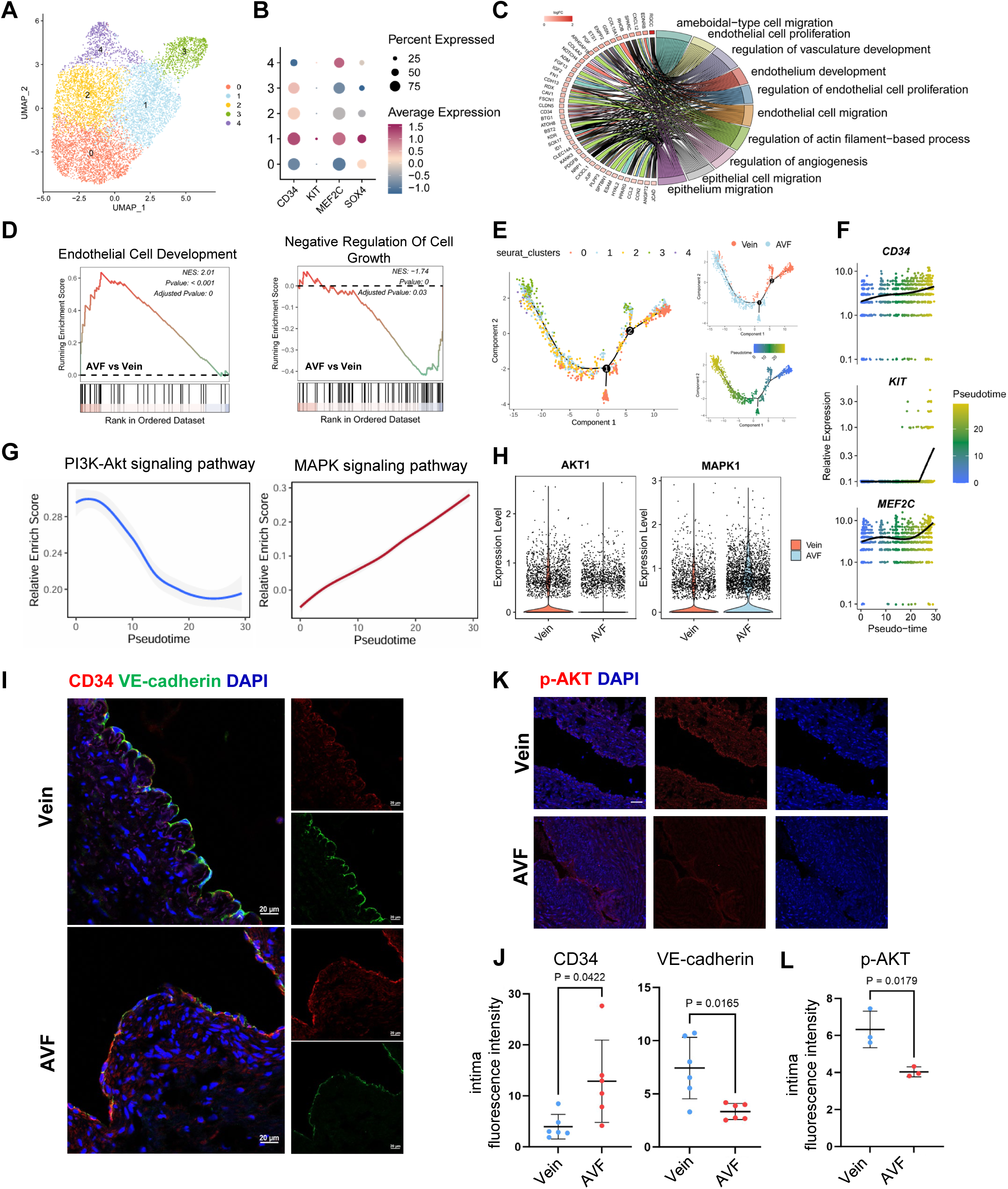
CD34 expression is increased whereas AKT signaling is attenuated in the endothelium of human AVF. **(A)** UMAP plot showing 5 distinct clusters of ECs in human normal vein and AVF. **(B)** Dot plots showing the expression levels of stemness-associated genes. **(C)** chord plot showing top enriched gene oncology functions in EC subcluster 1. **(D)** GSEA analysis showing the functional enrichment of AVF comparing to normal vein in EC subcluster 1. **(E)** Pseudo-time trajectory of EC subclusters with DDRTree algorithm for dimension reduction, along with the group and pseudotime score. **(F)** Trend line showing the expression changes of *CD34*, *KIT* and *MEF2C* along the pseudotime trajectory. **(G)** Trend line showing enrichment score of AKT and MAPK signaling along the pseudotime trajectory. **(H)** Violin plot showing *AKT1* and *MAPK1* expression level in the 4 EC in normal vein and AVF. **(I, J)** Immunofluorescence images of CD34 (red) and VE-Cadherin (green) in the AVF, with comparisons of fluorescence intensity per vessels. Scale bar=20 μm. **(K, L)** Immunofluorescence images of p-AKT (red) in the AVF, with comparisons of fluorescence intensity of p-AKT per vessels. Scale bar=50 μm. Statistical comparisons were performed using the two-tailed unpaired Welch’s t-test (J) and the two-tailed unpaired t-test (L).

To further investigate the phenotypic changes of this endothelial progenitor-like subcluster between AVF and normal veins, we performed GSEA and found that this subcluster was enriched for “Endothelial Cell Development” in AVF, whereas in normal vein controls, it was enriched for “Negative Regulation of Cell Growth” (Figure 7D). These findings are consistent with our observations in AVF mouse model. Subsequently, we used pseudotime analysis to simulate the dynamic phenotypic transition of normal veins following AVF surgery and observed a gradual upregulation of stemness-associated genes including *CD34*, *KIT*, and *MEF2C* (Figure 7E, 7F). Concurrently, pseudotime trajectory evaluation showed that AKT signaling progressively declined following AVF surgery, while MAPK signaling enrichment gradually increased (Figure 7G). The expression patterns of core genes *AKT1* and *MAPK1* were consistent with these pathway-level changes (Figure 7H). Immunofluorescence staining validated increased CD34 expression and decreased VE-Cadherin expression in human AVF tissues (Figure 7I, 7J). Consistent with AVF mouse model and in vitro shear stress experiments, decreased AKT phosphorylation and increased FoxO1 expression were also observed in human AVF tissues compared to control veins (Figure 7K,7L&S5E).

### Pharmacological Activation of AKT by SC79 Promotes the Differentiation of CD34^+^ Cells into Mature ECs in AVF Model

Based on our data showing critical role of AKT signaling in mediating maturation of CD34^high^ HUVECs, we propose that in vivo administration of SC79 would generate beneficial effect by reducing neointimal hyperplasia in AVFs. To evaluate the in vivo therapeutic efficacy of SC79, we administered SC79 or vehicle via repeated intraperitoneal injection to CD34-CreERT2; R26-tdTomato mice during AVF induction (Figure 8A). Four weeks later, we observed that AVF neointimal hyperplasia was significantly reduced in SC79-treated mice compared to vehicle-treated controls (Figure 8B). *En face* immunofluorescence staining showed that ECs in SC79-treated AVFs exhibited a more spindle-like morphology, resembling that of mature endothelium, accompanied by a marked decrease in CD34-derived tdTomato^+^ cells (Figure 8C,8D). Our data also showed that endothelial expression levels of VE-Cadherin and Claudin-5 in SC79-treated AVFs were significantly higher than that in the control group (Figure 8E, 8F). Together, these results confirm that SC79 promotes the mature differentiation of CD34^+^-derived ECs, suggesting its therapeutic promise to restore endothelial homeostasis and interrupt the vicious cycle of endothelial injury, maladaptive repair, and recurrent damage.

**Figure 8:**
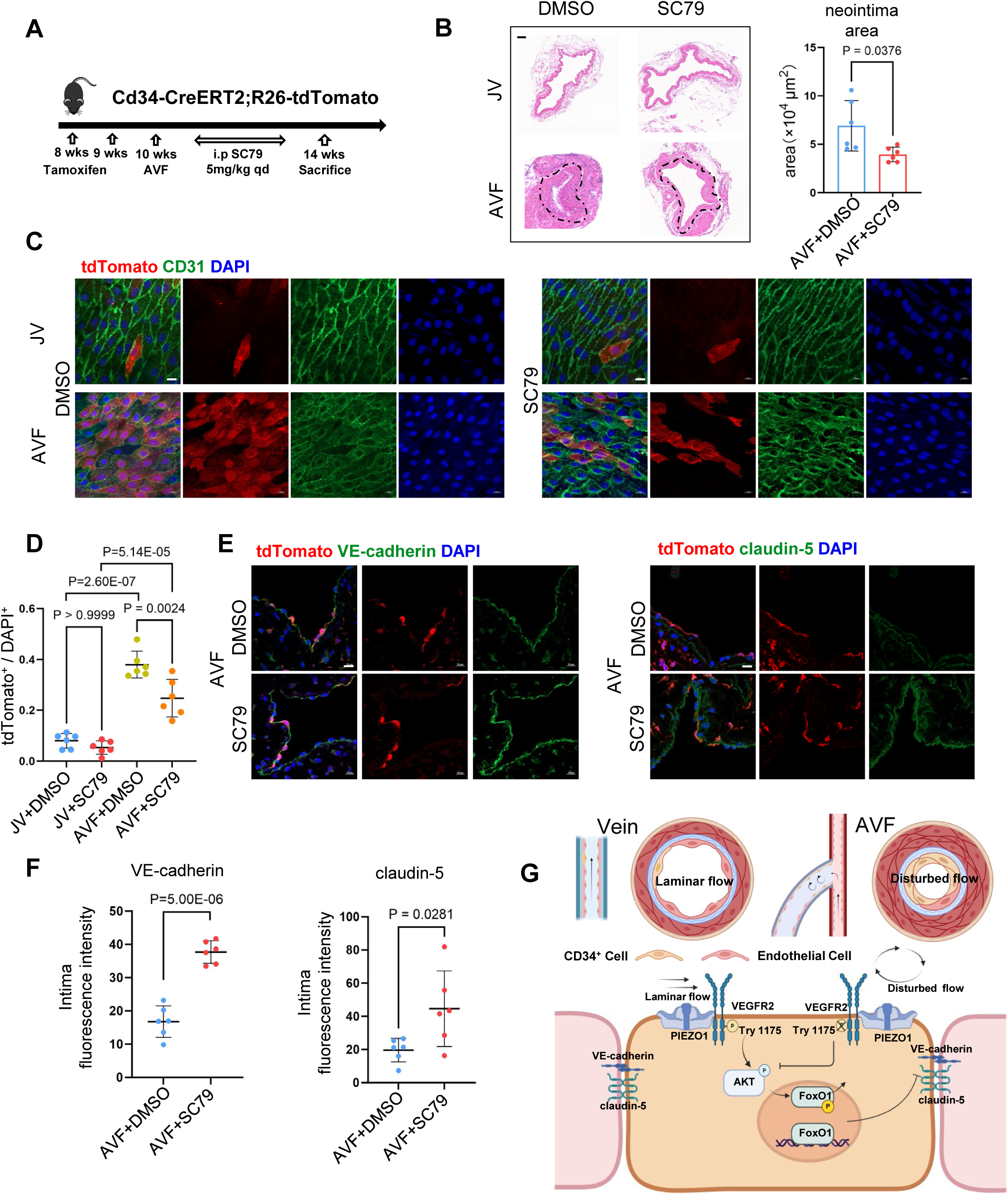
SC79 improved endothelial repair mediated by CD34^+^-derived cells in AVF. **(A)** Schematic diagram illustrating the construction strategy of CD34-CreERT2; R26-tdTomato mice with intraperitoneal injection of SC79 or vehicle. **(B)** Representative H&E staining images of AVF and jugular vein from Cd34-CreERT2; R26-tdTomato mice with intraperitoneal injection of SC79 or vehicle, with quantifications of neointima areas. **(C)** *En face* immunofluorescence images of tdTomato (red) and CD31 (green) in AVF and contralateral jugular vein from mice with intraperitoneal injection of SC79 or vehicle. Scale bar: 10 μm. **(D)** Dot plot graph showing the ratio of tdTomato^+^ cells to DAPI. **(E-F)** Immunofluorescence images of tdTomato (red), VE-Cadherin (green, left) and Claudin-5 (green, right) in AVF from mice with intraperitoneal injection of SC79 or vehicle, Scale bar: 10 μm. **(G)** Graphic abstract showing the role of PIEZO1-VEGFR2 complex and its downstream AKT-FoxO1 axis in regulating shear stress-induced differentiation of CD34^+^ cells into mature endothelium in AVF. Statistical comparisons were performed using the two-tailed unpaired Welch’s t-test (B; Claudin-5 in panel F), the two-tailed unpaired t-test (VE-Cadherin in panel F), and two-way ANOVA with Bonferroni correction (D).

## Discussion

The recruitment of CD34^+^ cells to endothelial injury sites has been recognized as an important process for maintaining normal endothelium function.^10,36^ Acute injuries caused by wire-induced denudation or transplantation primarily lead to the destruction of pre-existing endothelial cells (ECs) ^10,37,38^, thereby mainly triggering reparative differentiation of CD34^+^ cells to become mature endothelium. However, these models do not fully recapitulate the pathological features of arteriovenous fistulas (AVFs), where resident venous ECs exposed to abrupt and sustained increase in hemodynamic force after fistula creation. In such hemodynamic setting, endothelial injury is greater and persistently affected by abnormal shear stress, likely resulting in maladaptive differentiation of recruited CD34^+^ cells. In the present study, we demonstrate that persistent endothelial injury caused by disturbed flow in AVFs recruits abundant CD34^+^ cells to the injury site. The repair process at the injury site is disrupted by the abnormal hemodynamic environment, resulting in a repeated vicious cycle of injury-repair (injury, attempted repair, recurrent injury). These findings highlight an important knowledge gap in understanding how shear stress regulates the differentiation of CD34^+^ cells into mature endothelium. Elucidating the underlying mechanism of this process therefore is critical for understanding AVF failure and developing strategies to achieve functional endothelial regeneration in AVFs.

We sorted CD34^high^ and CD34^low^ HUVECs using flow cytometry and observed that their distinct responses to different shear stress patterns. Under laminar shear stress (LSS), CD34 expression was significantly downregulated, whereas the expression of endothelial maturation markers VE-cadherin and claudin-5 was markedly increased, indicating that LSS promotes the differentiation of CD34^high^ ECs into mature endothelium with enhanced junctional integrity. This finding is consistent with previous observations made from wire-injury models^10^, where CD34^+^ cells were capable of differentiating into mature endothelium and mitigating neointimal hyperplasia. In contrast, oscillatory shear stress (OSS) prevents the complete differentiation of CD34^high^ ECs into mature endothelium. This phenotype resembles the pathological features observed in AVF model, where substantial amount of CD34^+^ cells is recruited for endothelial repair, but functionally mature endothelium fails to establish and neointimal hyperplasia still occurs. Based on these findings, we propose that targeting specific flow-responsive signaling in CD34^+^ cells may represent a novel strategy to improve endothelial repair and reduce neointimal hyperplasia in AVFs.

Our bioinformatic analysis identified the mechanosensitive ion channel PIEZO1 as a pivotal mechanosensor to mediate shear stress-induced effect on CD34^+^ cell maturation. Prior studies show that PIEZO1 plays a critical role in the embryonic and postnatal endothelial development and homeostasis.^17,18^ However, little is known about whether PIEZO1-mediated mechanotransduction plays a key role in CD34^+^ cell differentiation in AVF. We knocked down PIEZO1 expression in CD34^high^ HUVECs and observed that PIEZO1 is essential for the differentiation of CD34^high^ HUVECs into mature endothelium induced by LSS and maintenance of stemness mediated by OSS as well. To consolidate these in vitro findings, we generated CD34-Cre^ERT2^; R26-tdTomato; Piezo1^flox/flox^ triple-transgenic mice to interrogate functional role of PIEZO1 in AVF. In PIEZO1-CKO group, we observed CD34-derived ECs exhibited a more mature endothelial repair and the neointimal hyperplasia was alleviated. These findings revealed that disturbed flow-activated PIEZO1 is a key negative regulator of the differentiation of CD34^+^ cells into mature ECs in AVFs.

Interestingly, we found that PIEZO1 forms a shear stress-sensing complex with VEGFR2 and selectively regulates downstream AKT but not ERK1/2 activation through shear stress-dependent VEGFR2 endocytic trafficking. The PIEZO1-VEGFR2 complex promotes phosphorylation of VEGFR2 at site 1175 and activates AKT under LSS, but it inhibits phosphorylation of VEGFR2 and AKT under OSS. Pharmacological blockade of AKT signaling impaired LSS-induced differentiation and maturation of CD34^high^ HUVECs. Conversely, pharmacological activation of AKT signaling promoted the differentiation and maturation of CD34^high^ HUVECs under OSS. Our findings showing PIEZO1-VEGFR2 interaction as a mediator of shear signal transmission extend further understanding of prior findings that report shear stress-mediated VEGFR2 regulation ^39–41^ and uncover a novel role of VEGFR2-AKT signaling in regulating shear stress-induced maturation of CD34^+^ cells. Importantly, in vivo pharmacological activation of AKT promoted the mature differentiation of CD34^+^-derived ECs and mitigated neointimal hyperplasia in AVFs, underscoring its therapeutic promise for restoring endothelial homeostasis.

Next, we explored the mechanism of shear stress-induced effect on maturation of endothelial junctions at transcriptional level. Bioinformatics analysis identified FoxO1, an important transcription factor downstream of AKT signaling^33,35,42,43^, is significantly upregulated in AVF. Given an established regulatory relationship among FoxO1, VE-cadherin and claudin-5, it is known that VE-Cadherin regulates claudin-5 expression via FoxO1 in EC^44,45^. In the present study, we examined whether FoxO1 directly controls gene expression program of junctional maturation in CD34^high^ HUVECs. FoxO1 overexpression markedly downregulated expression of VE-cadherin and claudin-5, which aligns with the reduced expression of VE-cadherin and claudin-5 observed in AVFs. Moreover, CUT& Tag analysis confirmed that FoxO1 directly binds to promoter region of VE-cadherin gene to inhibit its expression, supporting a transcriptional repressive effect of FoxO1 on endothelial adheren junction. Importantly, FoxO1 has previously been shown to regulate shear stress-induced proinflammatory gene expression, ^46,47^. Our data extend this concept by identifying endothelial junction disruption as a potential upstream mechanism linking FoxO1 activation to vascular inflammation. By transcriptionally repressing VE-cadherin and claudin-5, FoxO1 may thereby increase vascular permeability and facilitate immune cell infiltration into the remodeling vessel wall in AVF. Thus, FoxO1 may promote vascular hyperplasia in AVFs by coupling endothelial inflammatory response with defective junctional barrier. This mechanism provides a deeper explanation for how Akt activation contributes to attenuated neointimal hyperplasia in AVFs through FoxO1 inhibition.

Although our findings demonstrate that CD34^+^ cells primarily contribute to endothelial lineage differentiation, a subset of CD34^+^ cells in the AVF model may also differentiate into inflammatory cells and fibroblasts because of their multilineage differentiation potential^9–12^. Therefore, PIEZO1 deficiency in inflammatory cells and fibroblast lineages in PIEZO1-CKO mice may partially contribute to surgery-induced inflammation and fibrosis in AVF. However, our data show that CD34^+^ cells differentiate into more mature endothelium in PIEZO1-CKO mice, and it is well-known that matured endothelium protects against neointimal hyperplasia^48^. Moreover, in the AVF model, AKT activation effectively improved CD34^+^ cell-mediated endothelial repair, but the number of CD34-derived cells remained higher compared to normal veins, which was less pronounced than that observed in PIEZO1-CKO mice, indicating that other PIEZO1 downstream mechanisms may also contribute to endothelial repair. Additionally, while we identified an important role for the PIEZO1-VEGFR2 complex in shear stress sensing by CD34^high^ ECs, the exact binding mechanism requires further study and pharmacological target to this complex is currently unavailable to block VEGFR2-dependent shear stress signaling, warranting further investigation.

In summary, we have demonstrated that PIEZO1 plays a critical role in the differentiation and maturation of CD34^+^ cells during the repair of endothelial injury caused by abnormal shear stress in AVFs (Figure 8G). By forming a complex with VEGFR2, PIEZO1 transduces biomechanical cues to downstream AKT-FoxO1 axis, thereby regulating the shear stress-controlled differentiation of CD34^+^ cells into mature endothelium. Pharmacological activation of AKT improved the maturation of CD34^+^-derived ECs and alleviated neointimal hyperplasia in AVFs, providing a promising therapeutic strategy to restore vascular homeostasis.

## Nonstandard Abbreviations and Acronyms

AVF: arteriovenous fistula
AKT: protein kinase B, PKB
CD34: cluster of differentiation 34
CKO: CD34 conditional knock out
EC: endothelial cell
FoxO1: forkhead box protein O1
HUVEC: human umbilical vein endothelial cell
JV: jugular vein
LSS: laminar shear stress
OSS: oscillatory shear stress
PIEZO1: Piezo-type mechanosensitive ion channel component 1
SCR: scramble
VEGFR2: vascular endothelial growth receptor-2

## References

1. Cheung AK, Imrey PB, Alpers CE, Robbin ML, Radeva M, Larive B, Shiu Y-T, Allon M, Dember LM, Greene T, et al. Intimal Hyperplasia, Stenosis, and Arteriovenous Fistula Maturation Failure in the Hemodialysis Fistula Maturation Study. J. Am. Soc. Nephrol. 2017;28:3005–3013.

2. Lok CE, Huber TS, Orchanian-Cheff A, Rajan DK. Arteriovenous Access for Hemodialysis: A Review. JAMA. 2024;331:1307–1317.

3. Roy-Chaudhury P, Sukhatme VP, Cheung AK. Hemodialysis vascular access dysfunction: a cellular and molecular viewpoint. J. Am. Soc. Nephrol. 2006;17:1112–27.

4. Huber TS, Berceli SA, Scali ST, Neal D, Anderson EM, Allon M, Cheung AK, Dember LM, Himmelfarb J, Roy-Chaudhury P, et al. Arteriovenous Fistula Maturation, Functional Patency, and Intervention Rates. JAMA Surg. 2021;156:1111–1118.

5. Roy-Chaudhury P, Sukhatme VP, Cheung AK. Hemodialysis vascular access dysfunction: a cellular and molecular viewpoint. J. Am. Soc. Nephrol. 2006;17:1112–27.

6. Kipshidze N, Dangas G, Tsapenko M, Moses J, Leon MB, Kutryk M, Serruys P. Role of the endothelium in modulating neointimal formation: vasculoprotective approaches to attenuate restenosis after percutaneous coronary interventions. J. Am. Coll. Cardiol. 2004;44:733–9.

7. Chiu J-J, Chien S. Effects of disturbed flow on vascular endothelium: pathophysiological basis and clinical perspectives. Physiol. Rev. 2011;91:327–87.

8. Malek AM, Alper SL, Izumo S. Hemodynamic shear stress and its role in atherosclerosis. JAMA. 1999;282:2035–42.

9. Wang X, Wang R, Jiang L, Xu Q, Guo X. Endothelial repair by stem and progenitor cells. J. Mol. Cell. Cardiol. 2022;163:133–146.

10. Jiang L, Chen T, Sun S, Wang R, Deng J, Lyu L, Wu H, Yang M, Pu X, Du L, et al. Nonbone Marrow CD34+ Cells Are Crucial for Endothelial Repair of Injured Artery. Circ. Res. 2021;129:e146–e165.

11. Wang T, Gong H, Ye G, Chen R, Sun S, Huang X, Zhang B, Jiang L, Zhang Y, Chen T, et al. Multiple pathways of CD34+ cell differentiation during embryogenesis. Cell Death Differ. 2026;

12. Pu X, Zhu P, Zhou X, He Y, Wu H, Du L, Gong H, Sun X, Chen T, Zhu J, et al. CD34+ cell atlas of main organs implicates its impact on fibrosis. Cell. Mol. Life Sci. 2022;79:576.

13. Werner N, Junk S, Laufs U, Link A, Walenta K, Bohm M, Nickenig G. Intravenous transfusion of endothelial progenitor cells reduces neointima formation after vascular injury. Circ. Res. 2003;93:e17–24.

14. Griese DP, Ehsan A, Melo LG, Kong D, Zhang L, Mann MJ, Pratt RE, Mulligan RC, Dzau VJ. Isolation and transplantation of autologous circulating endothelial cells into denuded vessels and prosthetic grafts: implications for cell-based vascular therapy. Circulation. 2003;108:2710–5.

15. Kipshidze N, Dangas G, Tsapenko M, Moses J, Leon MB, Kutryk M, Serruys P. Role of the endothelium in modulating neointimal formation: vasculoprotective approaches to attenuate restenosis after percutaneous coronary interventions. J. Am. Coll. Cardiol. 2004;44:733–9.

16. Nonomura K, Lukacs V, Sweet DT, Goddard LM, Kanie A, Whitwam T, Ranade SS, Fujimori T, Kahn ML, Patapoutian A. Mechanically activated ion channel PIEZO1 is required for lymphatic valve formation. Proc. Natl. Acad. Sci. U. S. A. 2018;115:12817–12822.

17. Li J, Hou B, Tumova S, Muraki K, Bruns A, Ludlow MJ, Sedo A, Hyman AJ, McKeown L, Young RS, et al. Piezo1 integration of vascular architecture with physiological force. Nature. 2014;515:279–282.

18. Douguet D, Patel A, Xu A, Vanhoutte PM, Honoré E. Piezo Ion Channels in Cardiovascular Mechanobiology. Trends Pharmacol. Sci. 2019;40:956–970.

19. Jiang L, Zhang Y, Lu D, Huang T, Yan K, Yang W, Gao J. Mechanosensitive Piezo1 channel activation promotes ventilator-induced lung injury via disruption of endothelial junctions in ARDS rats. Biochem. Biophys. Res. Commun. 2021;556:79–86.

20. Friedrich EE, Hong Z, Xiong S, Zhong M, Di A, Rehman J, Komarova YA, Malik AB. Endothelial cell Piezo1 mediates pressure-induced lung vascular hyperpermeability via disruption of adherens junctions. Proc. Natl. Acad. Sci. U. S. A. 2019;116:12980–12985.

21. Gavard J, Gutkind JS. VE-cadherin and claudin-5: it takes two to tango. Nat. Cell Biol. 2008;10:883–5.

22. Tzima E, Irani-Tehrani M, Kiosses WB, Dejana E, Schultz DA, Engelhardt B, Cao G, DeLisser H, Schwartz MA. A mechanosensory complex that mediates the endothelial cell response to fluid shear stress. Nature. 2005;437:426–31.

23. Xu Q. Disturbed flow-enhanced endothelial turnover in atherosclerosis. Trends Cardiovasc. Med. 2009;19:191–5.

24. Chen KD, Li YS, Kim M, Li S, Yuan S, Chien S, Shyy JY. Mechanotransduction in response to shear stress. Roles of receptor tyrosine kinases, integrins, and Shc. J. Biol. Chem. 1999;274:18393–400.

25. Simons M, Gordon E, Claesson-Welsh L. Mechanisms and regulation of endothelial VEGF receptor signalling. Nat. Rev. Mol. Cell Biol. 2016;17:611–25.

26. Lansman JB. Endothelial mechanosensors. Going with the flow. Nature. 1988;331:481–2.

27. Tamargo IA, Baek KI, Kim Y, Park C, Jo H. Flow-induced reprogramming of endothelial cells in atherosclerosis. Nat. Rev. Cardiol. 2023;20:738–753.

28. Davis MJ, Earley S, Li Y-S, Chien S. Vascular mechanotransduction. Physiol. Rev. 2023;103:1247–1421.

29. Jin Z-G, Ueba H, Tanimoto T, Lungu AO, Frame MD, Berk BC. Ligand-independent activation of vascular endothelial growth factor receptor 2 by fluid shear stress regulates activation of endothelial nitric oxide synthase. Circ. Res. 2003;93:354–63.

30. Tzima E, Irani-Tehrani M, Kiosses WB, Dejana E, Schultz DA, Engelhardt B, Cao G, DeLisser H, Schwartz MA. A mechanosensory complex that mediates the endothelial cell response to fluid shear stress. Nature. 2005;437:426–31.

31. Chuntharpursat-Bon E, Povstyan O V, Ludlow MJ, Carrier DJ, Debant M, Shi J, Gaunt HJ, Bauer CC, Curd A, Simon Futers T, et al. PIEZO1 and PECAM1 interact at cell-cell junctions and partner in endothelial force sensing. Commun. Biol. 2023;6:358.

32. Simons M, Gordon E, Claesson-Welsh L. Mechanisms and regulation of endothelial VEGF receptor signalling. Nat. Rev. Mol. Cell Biol. 2016;17:611–25.

33. Paik J-H, Kollipara R, Chu G, Ji H, Xiao Y, Ding Z, Miao L, Tothova Z, Horner JW, Carrasco DR, et al. FoxOs are lineage-restricted redundant tumor suppressors and regulate endothelial cell homeostasis. Cell. 2007;128:309–23.

34. Wilhelm K, Happel K, Eelen G, Schoors S, Oellerich MF, Lim R, Zimmermann B, Aspalter IM, Franco CA, Boettger T, et al. FOXO1 couples metabolic activity and growth state in the vascular endothelium. Nature. 2016;529:216–20.

35. Deng H, Zhang X, Wang Y, Joshi D, Tellides G, Schwartz MA. FOXO1 Integrates Endothelial Hemodynamic, Inflammatory, and Metabolic Pathways in Atherosclerosis. Circ. Res. 2026;138.

36. Chen D, Abrahams JM, Smith LM, McVey JH, Lechler RI, Dorling A. Regenerative repair after endoluminal injury in mice with specific antagonism of protease activated receptors on CD34+ vascular progenitors. Blood. 2008;111:4155–64.

37. Chen T, Sun X, Gong H, Chen M, Li Y, Zhang Y, Wang T, Huang X, Wen Z, Xue J, et al. Host CD34+ cells are replacing donor endothelium of transplanted heart. J. Heart Lung Transplant. 2023;42:1651–1665.

38. Chen R, Li X, Zhu P, Yi X, Wang T, Chen H, Huang X, Ye G, Jiang J, Ong M, et al. CD34+ cell-Derived Endothelial Cells Orchestrate Vascular and Immune Remodeling in the Transplanted Liver. J. Hepatol. 2026;

39. Jin Z-G, Ueba H, Tanimoto T, Lungu AO, Frame MD, Berk BC. Ligand-independent activation of vascular endothelial growth factor receptor 2 by fluid shear stress regulates activation of endothelial nitric oxide synthase. Circ. Res. 2003;93:354–63.

40. Guo D, Chien S, Shyy JY-J. Regulation of endothelial cell cycle by laminar versus oscillatory flow: distinct modes of interactions of AMP-activated protein kinase and Akt pathways. Circ. Res. 2007;100:564–71.

41. Zeng L, Xiao Q, Margariti A, Zhang Z, Zampetaki A, Patel S, Capogrossi MC, Hu Y, Xu Q. HDAC3 is crucial in shear- and VEGF-induced stem cell differentiation toward endothelial cells. J. Cell Biol. 2006;174:1059–69.

42. Fosbrink M, Niculescu F, Rus V, Shin ML, Rus H. C5b-9-induced endothelial cell proliferation and migration are dependent on Akt inactivation of forkhead transcription factor FOXO1. J. Biol. Chem. 2006;281:19009–18.

43. Milkiewicz M, Roudier E, Doyle JL, Trifonova A, Birot O, Haas TL. Identification of a mechanism underlying regulation of the anti-angiogenic forkhead transcription factor FoxO1 in cultured endothelial cells and ischemic muscle. Am. J. Pathol. 2011;178:935–44.

44. Taddei A, Giampietro C, Conti A, Orsenigo F, Breviario F, Pirazzoli V, Potente M, Daly C, Dimmeler S, Dejana E. Endothelial adherens junctions control tight junctions by VE-cadherin-mediated upregulation of claudin-5. Nat. Cell Biol. 2008;10:923–34.

45. Giampietro C, Taddei A, Corada M, Sarra-Ferraris GM, Alcalay M, Cavallaro U, Orsenigo F, Lampugnani MG, Dejana E. Overlapping and divergent signaling pathways of N-cadherin and VE-cadherin in endothelial cells. Blood. 2012;119:2159–70.

46. Deng H, Zhang X, Wang Y, Joshi D, Tellides G, Schwartz MA. FOXO1 Integrates Endothelial Hemodynamic, Inflammatory, and Metabolic Pathways in Atherosclerosis. Circ. Res. 2026;138:e327592.

47. Luo J-Y, Cheng CK, He L, Pu Y, Zhang Y, Lin X, Xu A, Lau CW, Tian XY, Ma RCW, et al. Endothelial UCP2 Is a Mechanosensitive Suppressor of Atherosclerosis. Circ. Res. 2022;131:424–441.

48. Xiao Z, Postma RJ, van Zonneveld AJ, van den Berg BM, Sol WMPJ, White NA, van de Stadt HJF, Mirza A, Wen J, Bijkerk R, et al. A bypass flow model to study endothelial cell mechanotransduction across diverse flow environments. Mater. Today Bio. 2024;27:101121.

49. Chen K, Guan Z, Jin H, Aihemaiti M, Mou R, Yu M, Hu Y, Jiang L, Wang X, Yang H, et al. BACE2-Induced Aberrant Lymphatic Network Aggravates the Local Inflammation in Arteriovenous Fistulas With Hyperphosphatemia. Adv. Sci. (Weinh). 2025;12:e09632.

50. Sun X, Wang T, Gong H, Qiu Y, Zhang Y, Chen M, Xue J, Ye G, Mou R, Teng P, et al. Circulating CD34+ Fibroblast Progenitors Engaged in Heart Fibrosis of Allograft. Circ. Res. 2026;138:e326558.

51. Genet G, Boyé K, Mathivet T, Ola R, Zhang F, Dubrac A, Li J, Genet N, Henrique Geraldo L, Benedetti L, et al. Endophilin-A2 dependent VEGFR2 endocytosis promotes sprouting angiogenesis. Nat. Commun. 2019;10:2350.

52. Dunn KW, Kamocka MM, McDonald JH. A practical guide to evaluating colocalization in biological microscopy. American Journal of Physiology-Cell Physiology. 2011;300:C723–C742.

53. Ni Z, Lyu L, Gong H, Du L, Wen Z, Jiang H, Yang H, Hu Y, Zhang B, Xu Q, et al. Multilineage commitment of Sca-1+ cells in reshaping vein grafts. Theranostics. 2023;13:2154–2175.

